# Poly-unsaturated Fatty Acids from *Thamnidium elegans* and *Mortierella alpina* Suppress Prostate Cancer Cells Proliferation and Migration

**DOI:** 10.1101/2024.06.05.597505

**Authors:** Georgios Kalampounias, Panagiotis Dritsas, Dimitris Karayannis, Theodosia Androutsopoulou, Chrysavgi Gardeli, Seraphim Papanikolaou, George Aggelis, Panagiotis Katsoris

## Abstract

*Thamnidium elegans* and *Mortierella alpina* are two oleaginous fungi that belong to Mucoromycota that synthesize polyunsaturated fatty acids which are credited with multiple health benefits and possible anticancer properties. These fungi were cultivated on culture media with glucose or glycerol as a carbon source. After extracting the lipids, we transformed them into fatty acid lithium salts (FALS), which are water-soluble and absorbable mammalian cells, including DU-145 and PC-3 cancer cells. The two cell lines, both long-established prostate cancer models, were treated with FALS and indicated increased susceptibility to the lipid derivatives. The viability and proliferation rates were significantly reduced, as well as their migratory capabilities, which were significantly impaired compared to olive-oil-derived FALS, which were used as a control substance. We conclude that the FALS derivatives of microbial lipids from these organisms exhibit anticancer effects by suppressing the proliferation and migration of human prostate cancer cell lines.

## 1 Introduction

Polyunsaturated fatty acids (PUFA) have attracted the interest of the biotechnology sector due to their beneficial properties for human health. PUFA are taken up by the cell and used as an energy source or, under certain circumstances, for functional and structural purposes. The incorporation of PUFA carrying *cis* double bonds in cell membranes is crucial for functionality since saturation is invertedly correlated to membrane fluidity (Mukerjee *et al*., 2021). Additionally, their absorbance by the cells, may increase the cell’s susceptibility to oxidative stress damage since the aliphatic chains containing double bonds act as substrates for reactive oxygen species (ROS)-generating lipid hydroperoxides. This phenomenon, also known as lipid peroxidation, propagates exponentially if not suppressed by the cell’s antioxidant mechanisms and results in excessive oxidative damage, loss of membrane integrity, and finally cell death (Ayala, Muñoz & Argüelles, 2014). Flooding the cells’ catabolic pathways with unsaturated lipids drastically affects their oxidative stress levels, damages vulnerable intracellular structures and biomolecules, and significantly induces cell-death-related pathways such as ferroptosis and apoptosis (Marmunti & Catalá, 2007; Ge *et al*., 2009).

Gamma-linolenic acid (gLNA, 18:3n-6), arachidonic acid (ARA, 20:4n-6) and di-homo-gamma-linolenic acid (DGLA, 20:3n-6) are three fatty acids belonging to the omega-6 PUFA known for their cytoprotective/neuroprotective, anti-inflammatory, and anticancer properties, justifying the ever-increasing interest in research and industry (Kapoor & Huang, 2006; Chang *et al*., 2010; Wang, Lin & Gu, 2012; Xu & Qian, 2014; Tallima & El Ridi, 2017; Wang *et al*., 2021; Pereira *et al*., 2024). They are assumed to act through modulation of the cell metabolism, generation of reactive oxygen species, alteration of the membranes’ fluidity, and interfering with the intracellular signal transduction cascades (Das, 2020; Ponnampalam, Sinclair & Holman, 2021; Tallima & El Ridi, 2023). These fatty acids are believed to possess a high cytotoxic potential compared to other members of the group (Colquhoun & Schumacher, 2001; Ge *et al*., 2009; Wang, Lin & Gu, 2012; Bae *et al*., 2020; Sarparast *et al*., 2023; Tallima & El Ridi, 2023). However, a significant limiting factor in their extensive study and adoption was their scarcity. Since most food oils that contain significant amounts of them are mostly exploited as nutritional ingredients, their price remained high due to their low abundance and the cost of isolation and purification. Recently, advances in microbial biotechnology have allowed the production of microbial oils containing PUFA of high pharmaceutical value using various species that belong to Mucoromycota, especially *Mortierella,* and *Thamnidium*. These species are known for their ability to grow on cost-effective substrates such as food- and agricultural-industry byproducts (glycerol, pomace, etc.) and accumulate lipids that contain fatty acids like gLNA, DGLA, and ARA in significant quantities, while the isolation and purification process is relatively simple (Bellou *et al*., 2012; Hao *et al*., 2015; Fazili *et al*., 2022). Additionally, the whole process can be scaled up to an industrial level to produce great amounts of food- or even pharmaceutical-grade microbial lipids with potential therapeutic benefits.

Similar approaches, exploiting single-cell oils as substances of high added value from a pharmaceutical perspective, have been made in the past, investigating the lipids’ anticancer and antimicrobial potential (Alakhras *et al*., 2015; Sayegh *et al*., 2016; Kalampounias *et al*., 2024). In these articles, the lipids were not administered as free fatty acids, as they are characterized by poor solubility; nonetheless, they were transformed into fatty acid lithium/potassium salts (FALS/FAPS). This form is a liquid, water-soluble compound that does not require further transformations to be assimilable (e.g. the formation of liposomes or micelles) and has already been found to cause cell death by causing oxidative damage, and inducting apoptosis (Alakhras *et al*., 2015; Sayegh *et al*., 2016; Kalampounias *et al*., 2024). Given the variety of different fatty acids that can be produced through fermentation, it is important to recognize both the molecules possessing increased medicinal interest as well as to invest in the optimization of the culture process to achieve better yields and use more cost-effective strategies. This investigation aimed to produce microbial oils from the fungi *Mortierella alpina* CBS 343.66 and *Thamnidium elegans* CCF 1465 containing ARA, DGLA, and gLNA, respectively, and examine their effects on the DU-145 and the PC-3 prostate cancer cell lines. The oils were administered as FALS in the cell cultures, and key cellular functions were monitored to investigate any potential anticancer role.

## 2 Materials and methods

### 2.1 Biological material and culture conditions

The fungi *Mortierella alpina* CBS 343.66 and *Thamnidium elegans* CCF 1465 were kept for maintenance purposes on potato dextrose agar (Condalab, Madrid, Spain) at 7 ± 1 ^◦^C, and master cultures were regularly sub-cultured. The growth medium of *M. alpina* CBS 343.66 contained as a carbon and energy source glucose at 30 g/L, while *T. elegans* CCF 1465 was provided with glycerol at 30 g/L. Both growth media were supplemented with minerals, and their full compositions are presented in Table 1. Both media were nitrogen-limited, with yeast extract at 2.0 g/L being the only nitrogen source, while in the case of *M. alpina*, it was also deployed as a source of magnesium and ferrum. Medium pH was 6.5 ± 0.5 after sterilization at 121°C for 20 min and remained practically stable during cultivation, due to the high buffer capacity of the media containing Na_2_HPO_4_ and KH_2_PO_4_.

**Table 1.** The composition of the two synthetic growth media that were used for the cultures performed in this study.

| Compound | Supplier | *Mortierella* | *Thamnidium* |
| --- | --- | --- | --- |
|  |  | *alpina* CBS | *elegans* CCF |
|  |  | 343.66 | 1465 |
| Concentration (g/L) |  |  |  |
| Glycerol<br>(99.0% analytical grade) | Sigma-Aldrich (Darmstadt, Germany) | - | 30.00 |
| Glucose | Applichem (Darmstadt, Germany) | 30.00 | - |
| Yeast Extract | Condalab (Madrid, Spain) | 2.00 | 2.00 |
| (NH<sub>4</sub>)<sub>2</sub>SO<sub>4</sub> | Sigma-Aldrich | - | 2.00 |
| KH<sub>2</sub>PO<sub>4</sub> | Applichem | 12.00 | 7.00 |
| Na<sub>2</sub>HPO<sub>4</sub> | Applichem | 12.00 | 2.50 |
| MgSO<sub>4</sub>.7H<sub>2</sub>O | Sigma-Aldrich | - | 1.50 |
| CuSO<sub>4</sub>.5H<sub>2</sub>O | BDH (Poole, England) | 0.0001 | - |
| Co(NO<sub>3</sub>).6H<sub>2</sub>O | Merck (Darmstadt, Germany) | 0.0001 | - |
| FeCl<sub>3</sub>.6H<sub>2</sub>O | BDH | 0.08 | 0.15 |
| ZnSO<sub>4</sub>.7H<sub>2</sub>O | Merck | 0.001 | 0.02 |
| MnSO<sub>4</sub>.5H<sub>2</sub>O | Honeywell (Charlotte, North Carolina, USA) | 0.0001 | 0.06 |
| CaCl<sub>2</sub>.H<sub>2</sub>O | PENTA (Prague, Czechia) | 0.10 | 0.15 |

Submerged cultures were performed in 250-mL Erlenmeyer flasks containing 50 mL of growth medium. Flasks were aseptically inoculated with 1 mL of spore suspension containing 10^6^ fungal spores in a laminar flow apparatus. Incubation was held for approximately 6 days in an orbital shaker (ZHICHENG ZHWY 211C, Shanghai, China), equipped with a ventilation system, at 28 ± 1°C and an agitation rate of 180 ± 5 rpm.

### 2.2 Biomass determination

Fungal mycelia were collected by vacuum filtration through Whatman No. 1 paper. The harvested biomass was washed with distilled water in triplicate and then dried at 80 ^◦^C until constant weight. Finally, total biomass (x, g/L) was gravimetrically determined.

### 2.3 Extraction and purification of mycelial lipids

Total mycelial lipids were extracted in chloroform: methanol 2:1 (vol/vol) (Fisher Chemical, Hampton, NH, USA) as described elsewhere (Folch, Lees & Sloane, 1957). The extracts were filtered through Whatman No. 1 paper and washed with a 0.88 % (wt/vol) KCl (Honeywell Charlotte, NC, USA) solution to remove non-lipid components. Subsequently, the samples were evaporated under a vacuum using a Rotavapor R-20 device (BUCHI, Flawil, Switzerland). The total cellular lipid quantities (L, g/L) were gravimetrically determined and expressed as a percentage in the respective dry cell mass (L/x%).

### 2.4 Fatty acid composition of cellular lipids

To determine the fatty acid composition of all the samples used, total lipids were converted into their fatty acid methyl esters (FAME) in a two-step reaction, according to the AFNOR method (AFNOR, 1984). The produced FAME were analyzed with a Gas Chromatography (GC) (Agilent 7890A device, Agilent Technologies, Shanghai, China) device. The GC device was equipped with a flame ionization detector at 280°C, using an HP-88 (J&W Scientific, Folsom, CF, USA) column (60 m × 0.32 mm) for the analyses. Carrier gas was helium at a flow rate of 1 mL/min and the analysis was run at 200 °C. Peaks of FAMEs were identified through comparison to authentic standards.

### 2.5 Preparation of fatty acid lithium salts (FALS) solution pH 7

Saponification in 10 mL KOH (Carlo Erba, Emmendingen, Germany) 1 N ethanol (Fisher Chemical, Hampton, NH, USA) solution (95%) under reflux for 1 h and 45 min of 1 g of lipids occurred for glycerides cleavage. Acidification of the mixture was held with 10 mL HCl (Sigma-Aldrich, Saint-Louis, MO, USA) 4 N solution, and the free FA were extracted with 5 mL hexane (Sigma-Aldrich, Saint-Louis, MO, USA) in triplicate. Following, the organic phase was washed with distilled water until neutrality (pH 7) and dried over Na_2_SO_4_ (Honeywell, Charlotte, NC, USA). The organic phase was removed under a vacuum.

For FALS preparation, 1 g of FFA was diluted in 10 mL ethanol/diethyl ether (1:1; Sigma-Aldrich, Saint-Louis, MO, USA) and then LiOH (Sigma-Aldrich, Saint-Louis, MO, USA)1 N was added until pH 9. Thereafter, the solvents were removed under vacuum at T = 50 °C, and H_3_PO_4_ (Honeywell, Charlotte, NC, USA) 0.2 N was progressively added to the soap solution under stirring, until neutrality. Finally, distilled water was added to a final volume of 10 mL, resulting in a 10% wt/vol fatty acids-lithium salt aqueous solution.

### 2.6 Cell lines and culture

The DU-145 and the PC-3 cell lines were from ATCC and were used as models of human prostate cancer. The cells were cultured in RPMI 1640 cell culture medium (Biowest, Nuaillé, France) supplemented with 10% vol/vol fetal bovine serum (FBS) (Biowest, Nuaillé, France) and 1% vol/vol penicillin-streptomycin (Biowest, Nuaillé, France) in a CO_2_ incubator at 37 °C. All cell culture expendables (100 mm dishes, 24-well microplates, 6-well microplates, Boyden-Transwell Chambers) were purchased from Greiner Bio-One (Kremsmünster, Austria).

### 2.7 Proliferation assays

Equal numbers of cells were seeded inside 24-well plates and left to attach and grow for 24 h. After this interval, the media were aspirated and replaced with fresh RPMI 1640, supplemented with 10% FBS and the designated concentrations of FALS. Following 48 h of incubation, the viable cells were fixed using a 4% formaldehyde solution and subsequently stained using a Crystal Violet solution. The cell-bound stain was eluted using a 30% acetic acid solution and the optical density (O.D.) of the Crystal Violet solution was determined using a spectrophotometer at 595 nm. The O.D. values were converted to cell numbers using standard curves as described in previous studies (Kalampounias *et al*., 2024).

### 2.8 Wound healing-scratch test assay

The wound healing assay was used to study the cells’ wound healing capability following treatment with FALS (Rodriguez, Wu & Guan, 2005). Cells were seeded inside 6-well plates and cultured until the formation of a full monolayer. Scratches were made using P200 pipette tips and the old medium and debris were aspirated. RPMI 1640 supplemented with 10% FBS and selected doses of FALS were added, and photographs of the wounds were taken. The cells were incubated for 72 h and every 24 h, photographs were taken using a camera mounted on an inverted microscope.

### 2.9 Boyden chamber assay

To assess migration, equal cell numbers were placed in 8 μm inserts containing serum-free RPMI 1640 medium and designated doses of FALS. The inserts were then submerged in 24-well plates containing serum-supplemented medium. The cells were left to migrate for 24 h, and subsequently, the non-migrated cells were scrapped away while the migrated cells were fixed, stained, photographed, and counted as described in previous studies from our team (Kalampounias *et al*., 2024).

### 2.10 Statistical analysis

All experiments were conducted in triplicate and the resulting values were analyzed using the Prism 8 software (GraphPad, La Jolla, CF, USA). The proliferation data were analyzed using built-in models in Prism 8 for IC_50_ determination and the results are presented as scatter plots with a model-calculated fitting line. The wound-healing photographs were analyzed using the Wound Healing Tool for ImageJ and the resulting values were analyzed in Prism 8 (Suarez-Arnedo *et al*., 2020). The Boyden/Transwell chamber photographs were analyzed using the Cell Counter plug-in for ImageJ and the data were plotted using Prism 8.

## 3 Results

### 3.1 Biomass production, lipid accumulation, and fatty acid composition

*M. alpina* CBS 343.66 and *T. elegans* CCF 1465 were selected for this study due to their ability to accumulate PUFA-containing lipids. *T. elegans* cultivated on glycerol produced a high amount of biomass (x = 12.0 g/L) which was rich in lipids (L/x% = 49.0%, wt/wt). The most dominant fatty acid produced was oleic (OA) (18:1n-9) (29.7%, wt/wt), followed by palmitic acid (PA) (16:0) (23.3 %, wt/wt) and linoleic acid (LA) (18:2n-6) (22.1 %, wt/wt). Most importantly, lipids contained gLNA (i.e., 19.4 %, wt/wt) in significant quantities as well, highlighting the suitability of *T. elegans* as a gLNA producer. On the other hand, *M. alpina* was cultivated in a mineral medium with glucose as the sole carbon and energy source and produced 4.1 g/L of dry biomass which contained satisfying levels of cellular lipids (i.e., L/x% = 14.6%, wt/wt) (Table 2). The main cellular fatty acids produced were 16:0, representing 32.2% of total FA, followed by gondoic (20:1n-9) (19.8%, wt/wt), and OA (18.0%, wt/wt) (Table 2). Regarding the important polyunsaturated fatty acids produced and accumulated by this fungus, considerable amounts of DGLA (20:3n-6) (4.5%, wt/wt), ARA (3.5%, wt/wt) and gLNA (2.9%, wt/wt) were recorded. Finally, a lower percentage of alpha-linolenic acid (aLNA) (18:3n-3) was found. The aforementioned account for a total of PUFA that exceeds 10.0% (wt/wt) of the fatty acids of total lipids. Lastly, the fatty acid compositions of the negative control (olive oil) and positive control (eicosapentaenoic acid, EPA 20:5n-3, concentrate in the commercial form of E-EPA) of the following experiments are also presented in Table 2.

**Table 2.** Fatty acid composition of the methyl ester mixtures used in the fatty acid lithium salts. Experiments were performed in duplicate.

| Source of lipid | x<br>(g/L) | L/x%<br>(wt/wt) | Fatty acid composition (% , wt/wt) |  |  |  |  |  |  |  |  |  |  |  |  | Others |
| --- | --- | --- | --- | --- | --- | --- | --- | --- | --- | --- | --- | --- | --- | --- | --- | --- |
|  |  |  | 14:0 | 16:0 | 16:1n-7 | 18:0 | 18:1n-9 | 18:2n-6 | 18:3n-6 | 18:3n-3 | 18:4n-3 | 20:1n-9 | 20:3n-6 | 20:4n-6 | 20:5n-3 |  |
| *Mortierella alpina* CBS 343.66 | 4.1 | 14.6 | 1.0 | 32.2 | 0.2 | 8.4 | 18.0 | 8.3 | 2.9 | 0.9 | ND | 19.8 | 4.5 | 3.5 | ND | 0.2 |
| *Thamnidium elegans* CCF 1465 | 12.0 | 49.0 | 5.5 | 23.3 | ND | ND | 29.7 | 22.1 | 19.4 | ND | ND | ND | ND | ND | ND | ND |
| Olive oil | - | - | ND | 12.0 | 2.0 | 2.6 | 74.5 | 7.6 | ND | 0.7 | ND | ND | ND | ND | ND | 0.6 |
| E-EPA | - | - | ND | 0.5 | 0.4 | 3.1 | 9.2 | 0.6 | 0.5 | 2.3 | 1.7 | 4.4 | ND | ND | 75 | 2.3 |

**Abbreviations:** ND: Not-Detected

### 3.2 Effects on cell viability and proliferation

Following a 48-hour exposure of DU-145 and PC-3 cells to various concentrations of FALS from *T. elegans*, *M. alpina*, EPA concentrate (in the commercial form of E-EPA, used as a positive control) (Colquhoun & Schumacher, 2001), and olive oil (as a negative control), the IC_50_ values were calculated (Figure 1, Table 3). Among the four FALS formulations, the EPA-concentrate FALS exhibited the most evident cytotoxic and anti-proliferative action, and olive-oil-derived FALS was the mildest. The microbial lipid FALS fell in between the two controls, justifying their use and indicating PUFA-related dose-dependent cytotoxicity. The *T. elegans* lipids, containing about 19.4% wt/wt gLNA and in total 41.5% unsaturated fatty acids, exhibited an IC_50_ of ∼60 μg/mL in both DU-145 cells and PC-3 cells following 48 hours of incubation (Figure 1). The cell morphology after treatment with FALS had significantly altered, as shown in the figures, with the formation of lipid droplets being evident, as well as visible signs of cell swelling, dark spots, and apoptotic characteristics (Figure 2). FALS from *Mortierella alpina* demonstrated more severe cytotoxic effects, which were credited to the presence of DGLA in the increased concentration of 4.5% wt/wt, and that of ARA in the concentration of 3.5% wt/wt (Wang, Lin & Gu, 2012; Tallima & El Ridi, 2023).

**Fig. 1.**
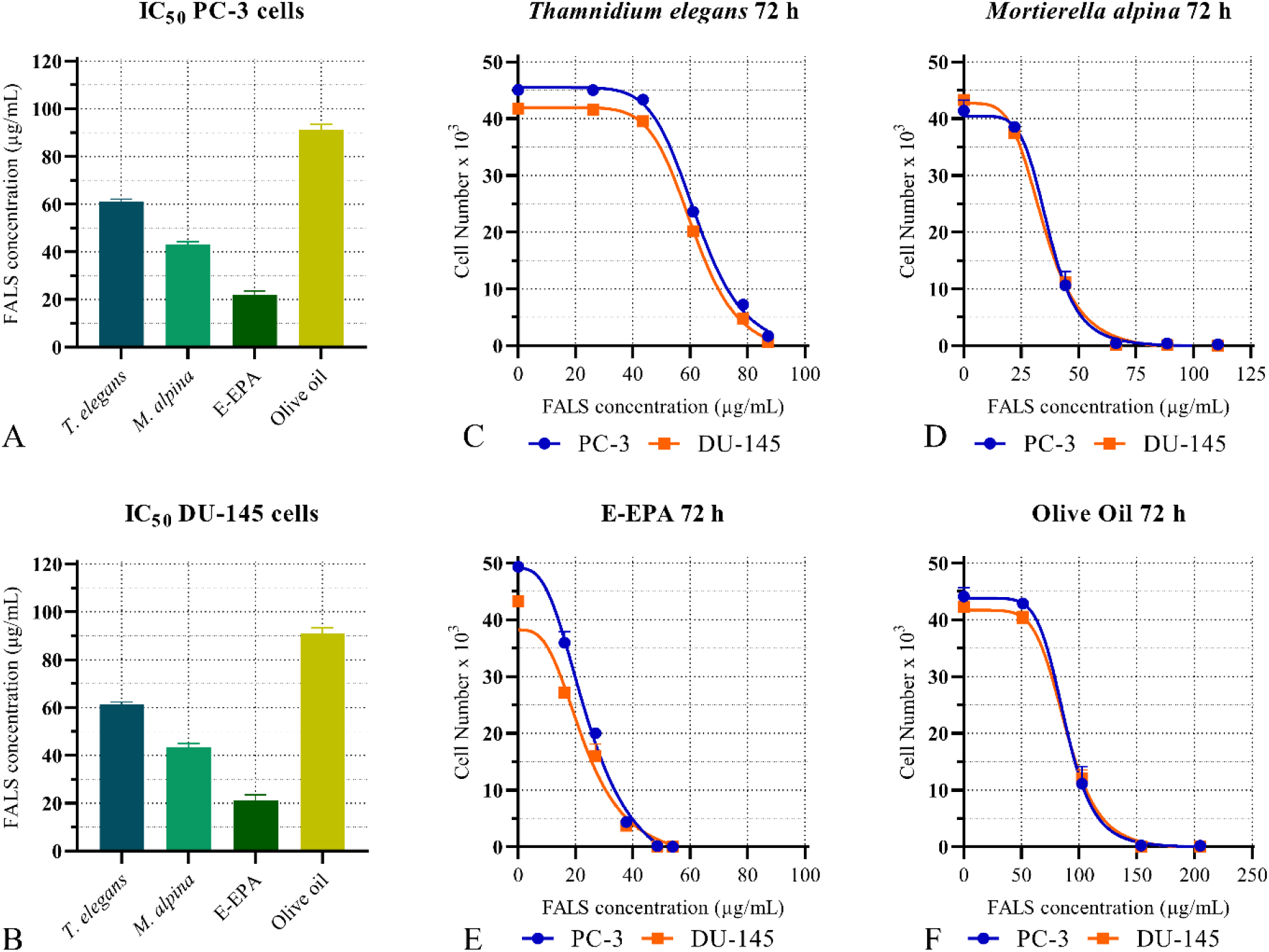
IC_50_ determination and proliferation curves of PC-3 and DU-145 cells. Following 48 h of incubation with increasing concentrations of FALS, the number of remaining alive cells was determined using the crystal violet assay. The data were analyzed using the built-in equations of the Prism 8 software. Each bar corresponds to the mean value of three repetitive IC_50_ calculations of the different lipid formulations ± the standard error of the mean (SEM). (A) PC-3 or (B) DU-145 cell IC50 values resulting from three independent experiments showing the different degrees of cytotoxicity between the tested FALS. The proliferation curves of PC-3 cells are annotated using the blue line and the blue dots, and those of DU-145 cells are indicated using the orange line and orange squares. Proliferation curve following incubation with FALS from: (C) *Thamnidium elegans*; (D) *Mortierella alpina*; (E) E-EPA; (F) Olive oil; All experiments were conducted in triplicate, and every point represents the average cell number ± standard error of the mean (SEM) values.

**Fig. 2.**
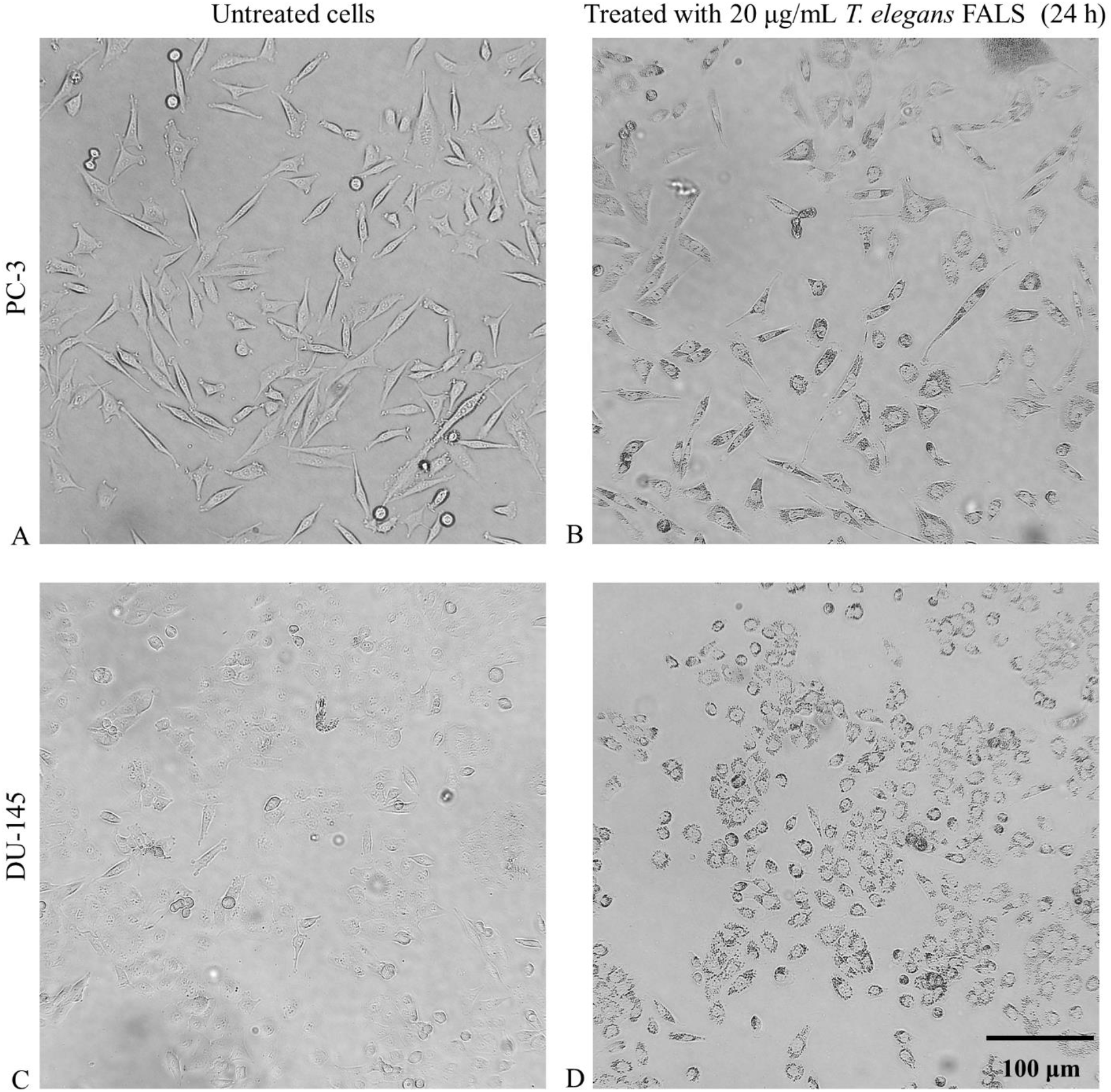
Cell morphology under a phase-contrast microscope. Following incubation with 20 μg/mL FALS for 24 h, cells grown on gLNAss coverslips were visualized under a phase contrast microscope. All photographs were taken at ×200 magnification. (A) Untreated PC-3 cells; (B) PC-3 treated with *T. elegans* FALS; (C) untreated DU-145 cells; (D) DU-145 cells treated with *T. elegans* FALS. Both cell lines exhibited darker coloration, increased and abnormal size, and visible granulation.

**Table 3.**
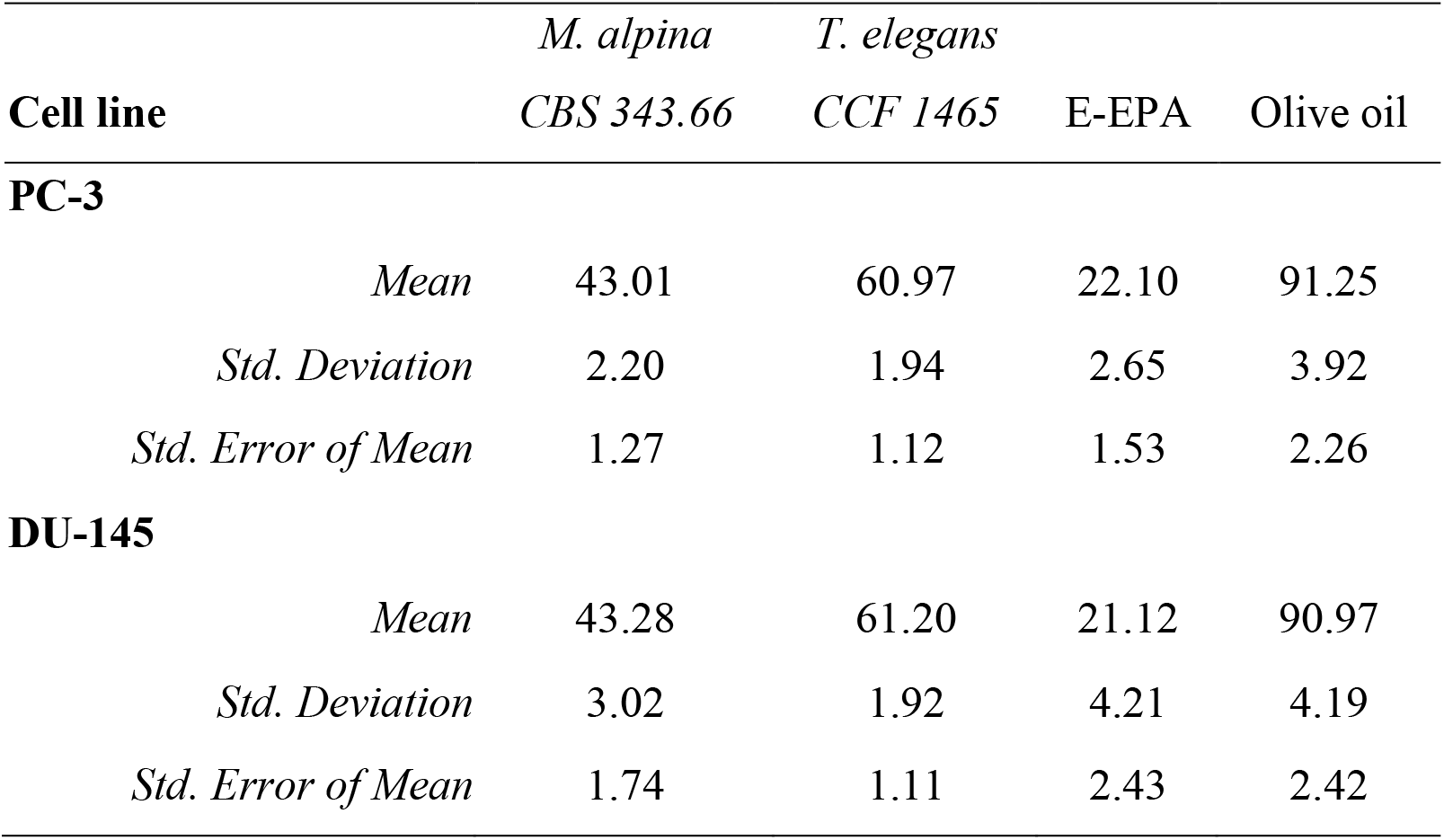
The IC_50_ value of PC-3 and DU-145 following 48 hours of incubation with a range of FALS concentrations.

|  |  | *M. alpina* | *T. elegans* |  |  |
| --- | --- | --- | --- | --- | --- |
| **Cell line** |  | *CBS 343.66* | *CCF 1465* | E-EPA | Olive oil |
| **PC-3** |  |  |  |  |  |
|  | *Mean* | 43.01 | 60.97 | 22.10 | 91.25 |
|  | *Std. Deviation* | 2.20 | 1.94 | 2.65 | 3.92 |
|  | *Std. Error of Mean* | 1.27 | 1.12 | 1.53 | 2.26 |
| **DU-145** |  |  |  |  |  |
|  | *Mean* | 43.28 | 61.20 | 21.12 | 90.97 |
|  | *Std. Deviation* | 3.02 | 1.92 | 4.21 | 4.19 |
|  | *Std. Error of Mean* | 1.74 | 1.11 | 2.43 | 2.42 |

### 3.3 Effects on wound healing ability

The effects of *T. elegans* and *M. alpina* on wound healing ability were also assessed. We performed dose-response experiments using various FALS concentrations (since every formulation had a significantly different IC_50_ value) and to compare them, we selected three main doses (10, 20, and 40 μg/mL) that were related to one another, and in this spectrum, differences regarding the FALS effects on cell migration could be documented (without significant cytotoxicity) (Figure 3). Even at the lowest concentration, the EPA-treated cell’s healing ability indicated significant impairment, justifying its selection as a positive control. The medium dose of 20 μg/mL led to significant migratory impairment by all formulations, with the most active being the EPA concentrate, followed by the *M. alpina* FALS and, to a lesser (but significant) point, the *T. elegans-*derived FALS. At this dose, the olive oil-derived FALS did not indicate significant wound healing impairment, also justifying the dose and control selection. Finally, the high dose of 40 μg/mL led to a greater effect by inhibiting the final wound closure, even within 72 h. As a control for the assay, cells treated only with 10% serum-containing medium were used which had a consistent pattern and rapid rate of wound healing.

**Fig. 3.**
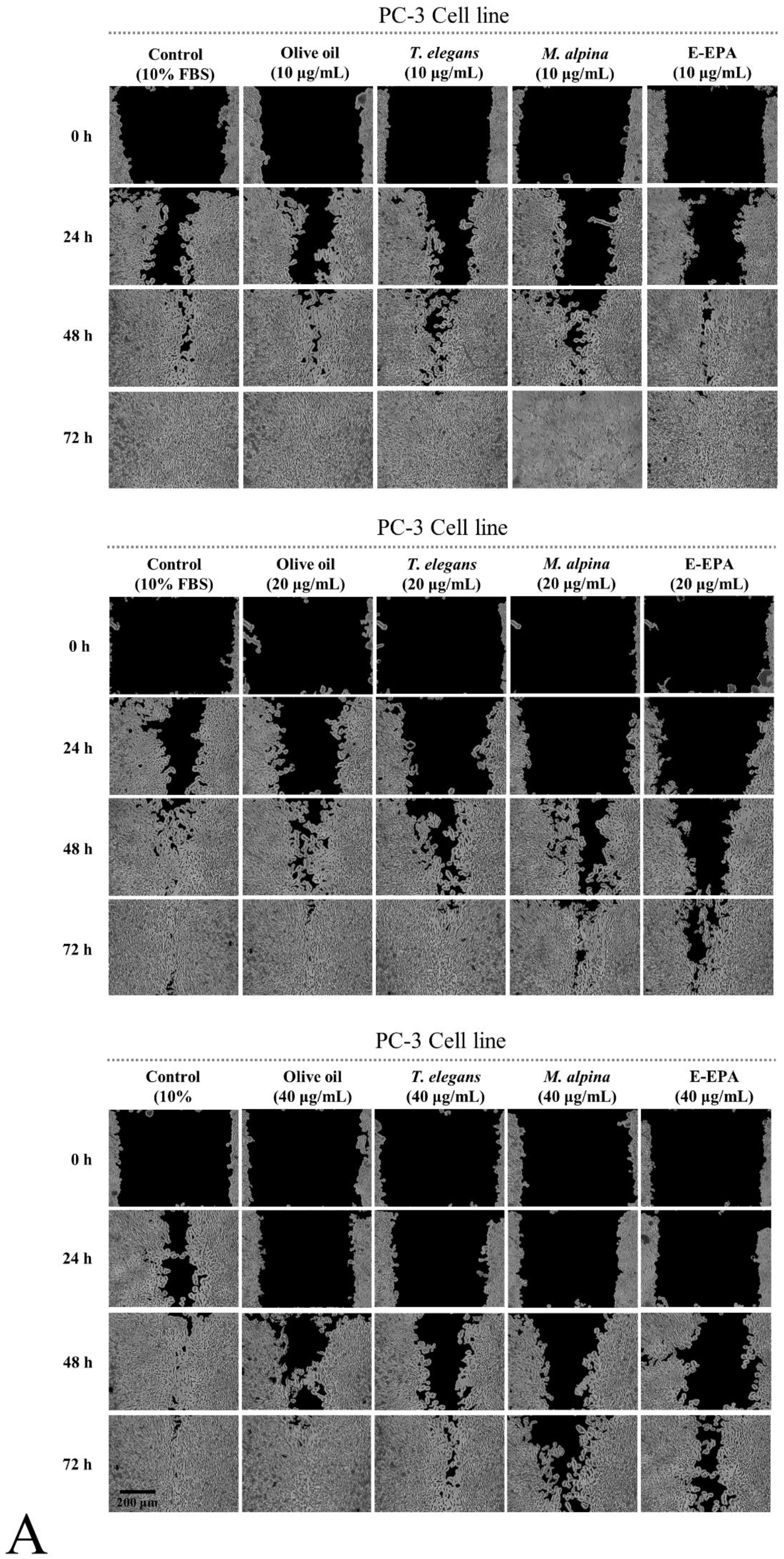
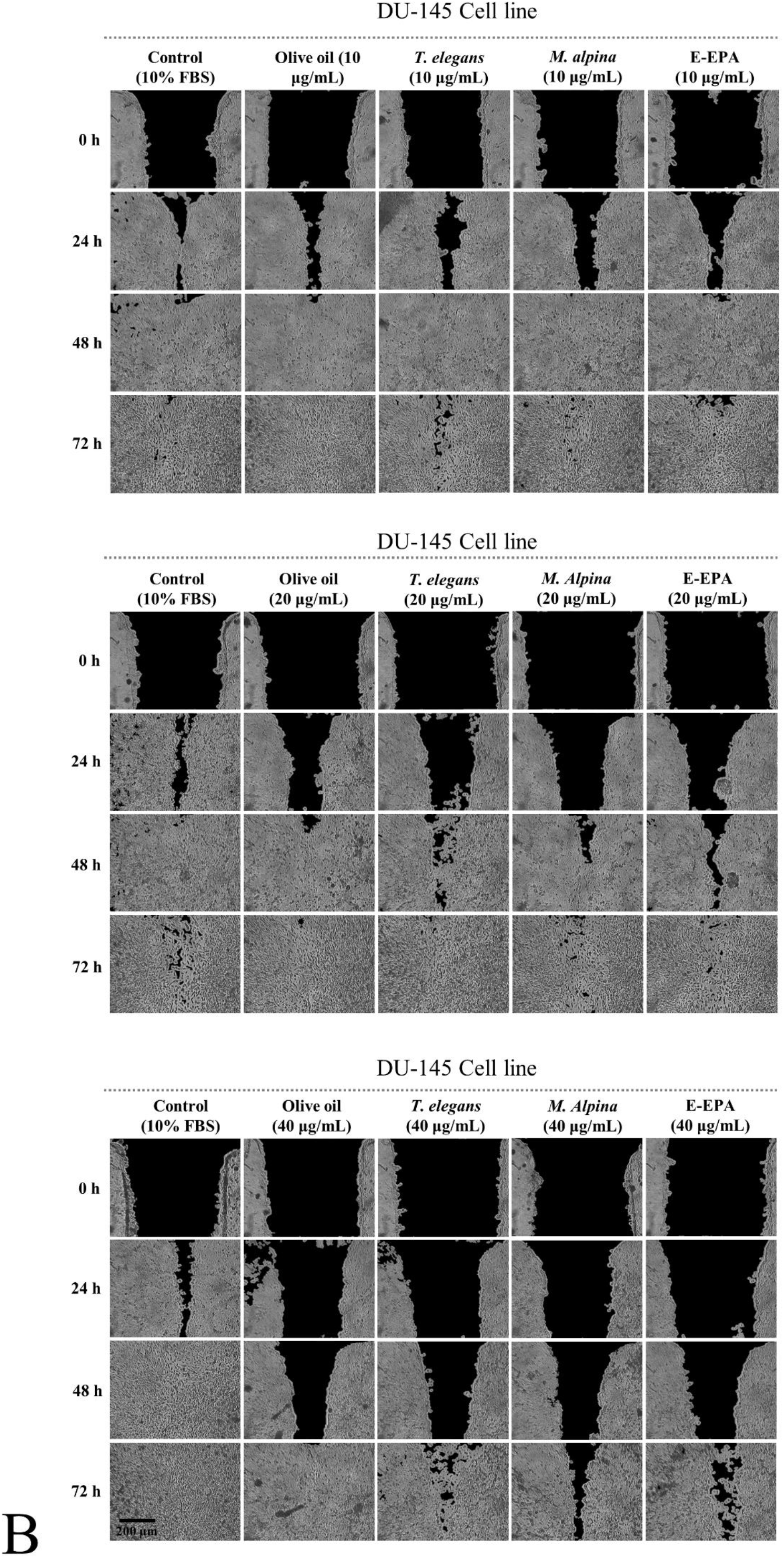
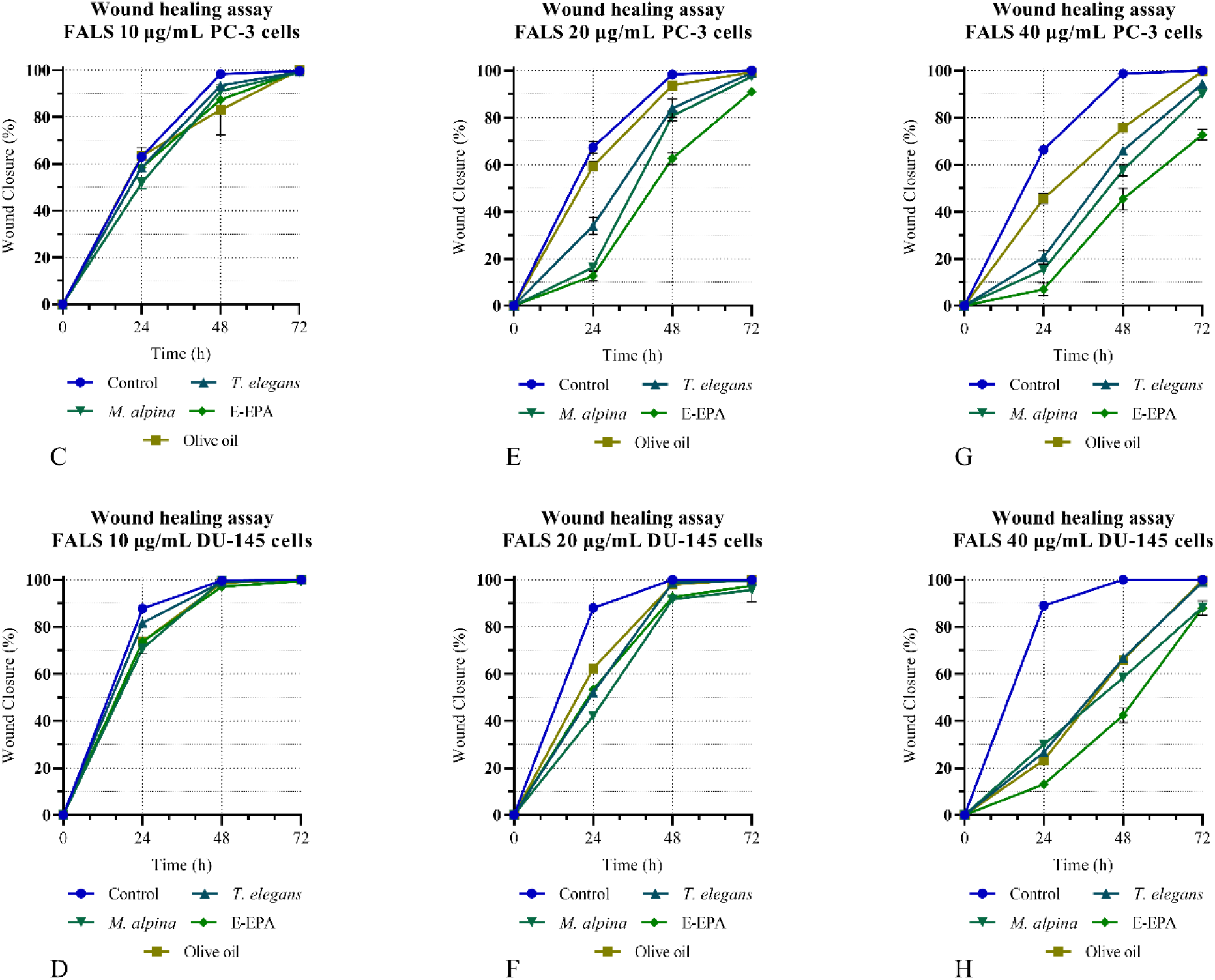
Wound healing assay. Using the scratch test, the effects of the various FALS formulations on the wound healing ability of (A) PC-3 and (B) DU-145 cells were assessed. Three different doses were administered (10, 20, and 40 μg/mL) after the wound formation, and photographs were taken at the key time points of 0, 24, 48, and 72 h. Cells supplemented only with 10% FBS were used as control samples. The data analysis was performed using the wound healing macro tool in the ImageJ software, and all the graphs were designed using Prism 8. The plots present the effects of the different doses on the two cell lines in the wound closure percentage during the key intervals. (C, D) PC-3 and DU-145 treated with 10 μg/mL FALS; (E, F) PC-3 and DU-145 treated with 20 μg/mL FALS; and (G, H) PC-3 and DU-145 treated with 40 μg/mL FALS; Each line represents a different FALS formulation, and each point is the mean percentage extracted from three photographs. The error bars correspond to the standard error of the mean (SEM).

### 3.4 Effects on cancer cell migration

Regarding migration, a similar experimental design to wound healing experiments was followed by treating PC-3 and DU-145 cells with a medium FALS concentration (20 μg/mL) for 24 h. This dose was selected using the wound healing experiments, where at 20 μg/mL: the olive-oil-treated cells had a similar response to untreated cells; the *T. elegans*-derived FALS impaired migration to a greater point; *M. alpina*-FALS imposed a more intense migration inhibition; and finally, the EPA-treated cells had the greater degree of migratory ability impairment. Regarding *T. elegans* FALS, treatment for 24 h caused a 40% decrease in the migration of PC-3 cells and a 50% decrease in that of DU-145 cells, respectively (Figure 4). *M. alpina* FALS reduced migration levels of PC-3 cells by 70% in PC-3 cells and by 75% in DU-145. The effects of EPA concentrate were even greater, causing an 80% reduction in cell migration, while the use of olive oil FALS at the same concentration only reduced migration by 40-50% in both cell lines, which was significantly lower compared to the microbial-derived FALS. These observations, emphasize the potential antitumor activity gLNA-, DGLA-, and ARA-rich lipids can exhibit and support the further use of lithium salts (FALS) as a way to administer them as has already been shown by previous studies (Alakhras *et al*., 2015; Sayegh *et al*., 2016; Kalampounias *et al*., 2024).

**Fig. 4.**
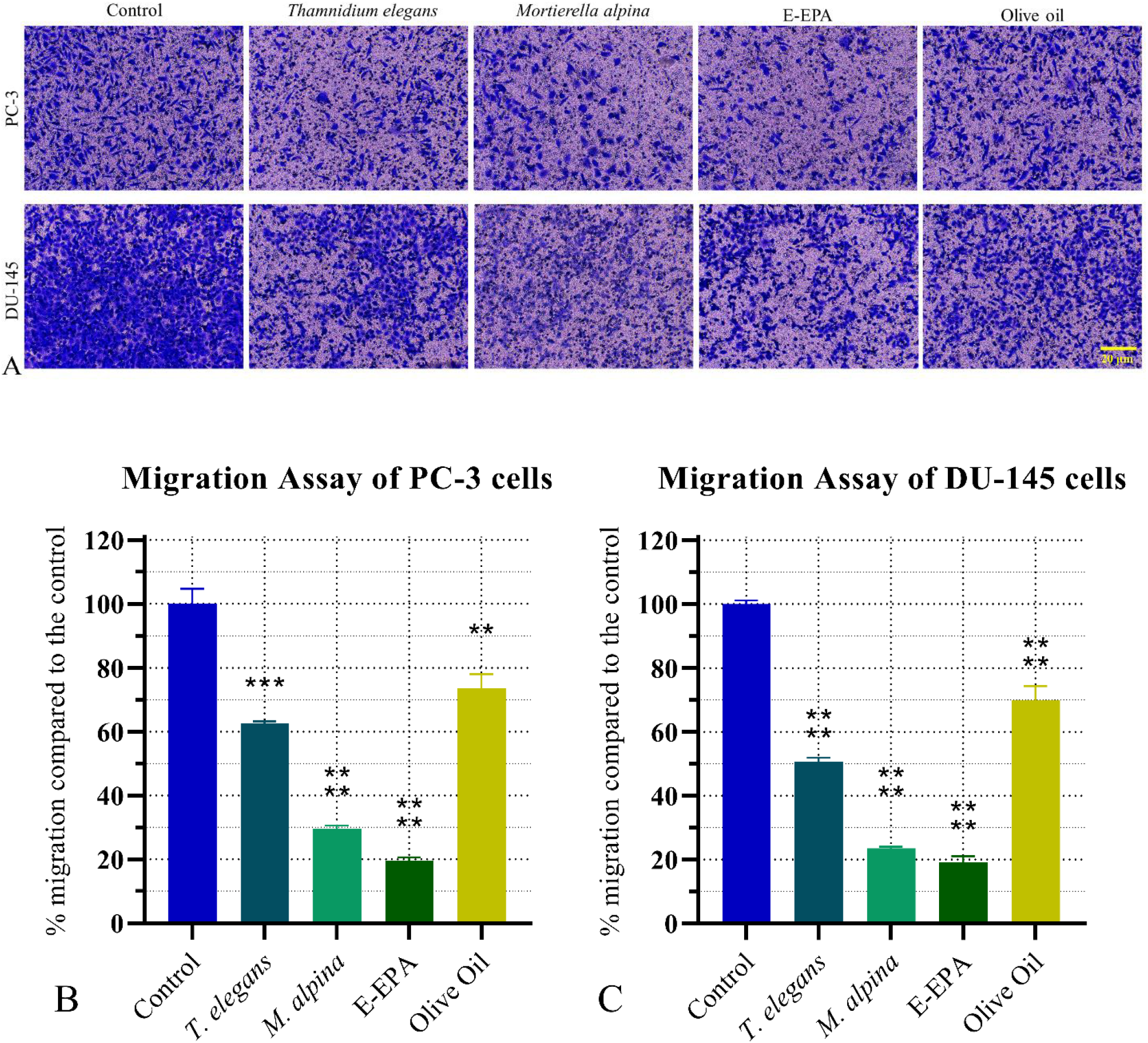
Migration assay. Using Boyden chambers, the effects of 20 μg/mL FALS on the migratory capability of PC-3 and DU-145 cells after a 24-hour incubation were assessed. To study the migration, FALS were present in the cell medium (inside the insert) while the chemoattractant medium (10% FBS did not contain FALS. (A) After 24 h, the cells crossing the filter were fixed, and stained with crystal violet, and photographs were taken using a photonic microscope. (B, C) The images were analyzed using the built-in tool cell counter by ImageJ. The data analysis was performed on Prism 8 and the migration rate of (B) PC-3, and (C) DU-145 cells is presented as mean percentages of migration compared to the control (no FALS). Three independent experiments were performed to extract the mean values and the error bars correspond to the standard error of the mean (SEM).

## 4 Discussion

In this study, lipids from *T. elegans* and *M. alpina* were administered for the first time to human prostate cancer cell lines, and an assessment of the cells’ ability to proliferate and migrate was performed. Lipids from both microorganisms were transformed into FALS, a bio-assimilable form of the insoluble long-chain PUFA that has previously been used for in vitro assays by both our research group (Alakhras *et al*., 2015; Sayegh *et al*., 2016; Kalampounias *et al*., 2024) and other researchers (Kairemo *et al*., 1998; Ilc *et al*., 1999). Lithium, in the form of lithium chloride and lithium carbonate, is a new promising agent in anticancer research since its effectiveness against brain tumors is under investigation (Natale *et al*., 2023). Therefore, FALS, by combining two potential tumor-suppressor substances that target cancer cells, might be a formulation of high pharmaceutical value that could be used as an adjunct therapy in the future due to its low toxicity compared to conventional chemotherapies.

*T. elegans* CCF 1465 cultivated on glycerol produced significant amounts of biomass, confirming herein that this fungus is an exception to the general rule that considers glycerol as a not very appropriate carbon source for Mucorales due to the low uptake rates that have been recorded (Bellou *et al*., 2012; Dritsas & Aggelis, 2023). More importantly, this fungus accumulated lipids rich in gLNA, a PUFA of increased significance regarding potential anticancer properties, which justified the selection of this strain. *In vitro* data have provided much evidence on the suppressive role of gLNA on cancer cell proliferation and migration of many cancer types, among which also lies prostate cancer (Kairemo *et al*., 1998; Ilc *et al*., 1999; Robbins *et al*., 1999; Meng *et al*., 2013; Alakhras *et al*., 2015; Kalampounias *et al*., 2024). Previous studies had shown that *T. elegans* CCF 1465 lipids in the form of fatty acid potassium salts (FAPS) exhibited anticancer properties against the breast cancer MCF-7 cell line (Sayegh *et al*., 2016), and the current study greatly expands and verifies these results since cytotoxicity against prostate cancer cells is demonstrated as well as inhibitory effects on cell migration. All these *in vitro* data as well as our study significantly support experiments conducted in 2006, where Pham et al. showed that incorporation of gLNA-rich foods in rat diets reduced the size of prostate tumors (Pham, Vang & Ziboh, 2006), a discovery that truly questions whether more research should be conducted in this direction. gLNA is not a common fatty acid in human nutrition, and it mainly reaches us in trace amounts, due to the limited organisms that are able to synthesize it. However, some oleaginous fungi produce it, and it is noteworthy that many of them can be grown in laboratory- and/or industrial-scale settings. Examples of such producers are *Cunninghamella echinulata*, *Cunninghamella elegans*, and *Thamnidium elegans*, all of which have been found to grow on glucose or even glycerol as a carbon source, and their fatty acids possess anticancer properties, as indicated by in vitro experiments. Overall, growing *T. elegans* on glycerol, which is a byproduct of the agricultural and food industries in combination with the successful accumulation of gLNA-rich lipids is a great example of a cost-effective method and substrate that can lead to the production of high-added value substances. On the other hand, *M. alpina* CBS 243.66 was selected due to its ability to synthesize gLNA, DGLA, and ARA. gLNA is a precursor to the synthesis of DGLA which is the intermediate of ARA biosynthesis (Nykiforuk *et al*., 2012). Intriguingly, Wynn and Ratledge considered the elongation of the gLNA to DGLA as the rate-limiting step for ARA production as, according to the authors, an enhancement of the elongase activity could lead to an increase in ARA production (Wynn & Ratledge, 2000). In general, DGLA is much less abundant in the human diet and it is present only in some seed oils, which are relatively uncommon which has been credited with strong tumor-suppressive properties following *in vitro* experiments on human cancer cells (Wang, Lin & Gu, 2012; Xu & Qian, 2014; Sarparast *et al*., 2023). ARA is a relatively common fatty acid, available in many meat and dairy foods, that may possess strong tumoricidal properties (Tallima & El Ridi, 2023). *M. alpina* CBS 243.66 might not produce large quantities of ARA, though, its content in ARA could be improved by growing the fungus at a lower temperature (i.e., T = 18 °C), as Lindberg and Molin (1993) proposed for this strain (Lindberg & Molin, 1993).

*M. alpina* FALS were found to be significantly more cytotoxic compared to *T. elegans* lipids, an observation that may be a result of the combined action of DGLA and ARA in the cancer cells. Tallima et al. in 2023 concluded that the cytotoxicity observed on numerous cell lines following incubation with ARA, was a result of changes in cell membrane, that possibly activate contact inhibition as well as the activation of cell surface membrane-associated neutral sphingomyelinase (nSMase) that led to an accumulation of Cer that ultimately suppressed the cells’ proliferation (Tallima & El Ridi, 2023). Bae et al. in 2020 also observed inhibitory effects of ARA on the human colon cancer cell line HT-29 (Bae *et al*., 2020). They proposed that the cells in the presence of ARA were unable to synthesize the necessary fatty acids needed for mitotic activity, as a result of an ARA-imposed inhibition of lipogenesis. An induction of apoptosis was also explained as a result of endoplasmic reticulum (ER) stress, which would be a direct consequence of the incorporation of vast quantities of fatty acids in the ER membranes (Bae *et al*., 2020). This observation is in accordance with the altered morphology of PC-3 and DU-145 cells we observed, both of which showed signs of lipid droplets formation inside their ER, which is a sign of increased lipid assimilation. The induction of apoptotic death has also been reported in the past in human chronic myeloid leukemia cells (Rizzo; *et al*., 1999), indicating a potential universal mode of action against multiple cancer types.

Moreover, DGLA, the other major PUFA found in *M. alpina* FALS, also contributes to the observed proliferation and migration suppression. DGLA has been reported to cause oxidative damage to cancer cells, thus leading to excessive peroxidation which ultimately provokes apoptosis and ferroptosis (Das & Madhavi, 2011; Wang, Lin & Gu, 2012; Xu & Qian, 2014; Sarparast *et al*., 2023). In general, at the biochemical level, the cytotoxic actions of DGLA are considered similar (if not equal) to those of gLNA. Both fatty acids are metabolic precursors of high importance in the class I prostaglandin pathway (PGE1). This class of prostaglandins and their various derivatives (e.g. leukotrienes), besides their hormone-like roles, are also credited with cytoprotective roles and decreases in thrombogenicity (Sinzinger *et al*., 1996). In 2020, Perez et al. showed that dietary consumption of DGLA by the nematode *Caenorhabditis elegans* led to germline cell ferroptosis. They also monitored the effects of DGLA administration on the human fibrosarcoma cell line HT-1080 and confirmed that ferroptosis was also being induced, leading to the cancer cells’ death.

All these data together indicate that the lipid formulations produced by selected Mucoromycota can possess significant anticancer properties and lead to significant suppression of tumor cells’ proliferation and migration. It is likely that all of the analyzed mechanisms perplex and combine to lead to tumoricidal effects; therefore, more research is warranted in order to extrapolate. FALS have been repeatedly used during the last 25 years as fatty acid carriers for in vitro studies, and given the positive outcomes those studies produce, it may be a good idea to employ more complex and accurate tumor models like organoids, 3D cultures, and ultimately organism models.

## 5 Conclusions

This study highlights the use of the filamentous fungi *T. elegans* CCF 1465 and *M. alpina* CBS 243.66 as producers of single cell oils (SCO) with high anticancer potential. After conversion to the drastic form of FALS, the effects of fatty acids derived from the above-mentioned microorganisms on cell proliferation and migration were studied, and significant inhibitory effects were documented regarding both parameters. These results fortify the use of FALS as a novel way to administer fatty acids to cells in in-vitro conditions and validate the suspected anticancer properties of gLNA, DGLA, and ARA encouraging further research regarding their mechanism of action and possible utilization. Additionally, the exploitation of industrial-grade glycerol as a substrate for *T. elegans* growth highlights the possibility of exploiting low-value carbon sources, often treated as byproducts or even waste, to produce pharmaceutical substances. *T. elegans* was found to accumulate slightly less cytotoxic PUFA compared to *M. alpina*, which mostly accumulated DGLA and ARA.

## Abbreviations

aLNA: alpha-linolenic acid
ARA: arachidonic acid
DGLA: di-homo-gamma-linolenic acid
EPA: eicosapentaenoic acid
ER: endoplasmic reticulum
FA: fatty acid
FAME: fatty acid methyl ester
FFA: free fatty acid
FALS: fatty acid lithium salts
FAPS: fatty acid potassium salts
gLNA: gamma-linolenic acid
LA: linolenic acid
ND: not detected
OA: oleic acid
PA: palmitic acid
PGE1: prostaglandin E1
PUFA: polyunsaturated Fatty Acids
ROS: reactive oxygen species
SCO: single cell oil

## Acknowledgments

The research within this article was financially supported by the project entitled “Biotransformation of glycerol into high pharmaceutical-value poly-unsaturated fatty acids (PUFAs)” (Acronym: Glycerol2PUFAs, project code HFRI-FM17-1839) financed by the Hellenic Foundation for Research & Innovation (H.F.R.I.), Nea Smyrni—Greece (action: “1st Call for H.F.R.I. Research Projects to Support Faculty Members & Researchers and Procure High-Value Research Equipment”).

## Authorship

Panagiotis Katsoris (P.K.), George Aggelis (G.A.), and Sepaphim Papanikolaou (S.P.) conceived and designed the study. George Kalampounias (G.K.), Panagiotis Dritsas (P.D.), Dimitris Karayannis (D.K.), Theodosia Androutsopoulou (T.A.), and Chrysavgi Gardeli (C.G.) provided the required methodology. G.K., P.D., D.K., and T.A. provided the appropriate software and performed the formal data analysis. G.K., P.D., and D.K. performed the experiments and their validation, were responsible for the data curation, and prepared the original draft. C.G., S.P., G.A., and P.K. provided the resources that made this work feasible. G.K and T.A. were responsible for the results’ visualization. P.K. supervised the project, and S.P. was responsible for administration and funding acquisition. S.P., G.A. and P.K. reviewed and edited the final manuscript. All authors contributed to the draft’s preparation, have read it, and agreed to the published version.

## Ethics statement

No human or animal subject were used in this research.

